# Lower Extremity Long-Latency Reflexes Differentiate Walking Function After Stroke

**DOI:** 10.1101/588111

**Authors:** Caitlin L. Banks, Virginia L. Little, Eric R. Walker, Carolynn Patten

## Abstract

The neural mechanisms of walking impairment after stroke are not well characterized. There is a need for a neurophysiologic marker that can unambiguously differentiate functional status and potential for walking recovery. The long-latency reflex (LLR) is a supraspinally-mediated response that integrates sensorimotor information during movement. It is hypothesized that lower extremity LLRs contribute to regulation of motor output during walking in healthy individuals. The goal of the present study was to assess the relationship between lower extremity LLRs, measures of supraspinal drive, and walking function. Thirteen individuals with chronic post-stroke hemiparesis and thirteen healthy controls performed both isometric and dynamic plantarflexion. Transcranial magnetic stimulation (TMS) assessed supraspinal drive to the tibialis anterior. LLR activity was assessed during dynamic voluntary plantarflexion and individuals post-stroke were classified as either LLR present (LLR+) or absent (LLR-). All healthy controls and nine individuals post-stroke exhibited LLRs, while four did not. LLR+ individuals revealed higher clinical scores, walking speeds, and greater ankle plantarflexor power during walking compared to LLR- individuals. LLR- individuals exhibited exaggerated responses to TMS during dynamic plantarflexion relative to healthy controls. This LLR- subset revealed dysfunctional modulation of stretch responses and antagonist supraspinal drive relative to healthy controls and the higher functioning LLR+ individuals post-stroke. These abnormal responses allow for unambiguous differentiation between individuals post-stroke and are associated with multiple measures of motor function. These findings provide an opportunity to distinguish among the heterogeneity of lower extremity motor impairments present following stroke by associating them with responses at the nervous system level.

## 1. Introduction

Walking is often impaired following stroke, putting individuals at high risk for falls (Jørgensen et al. 2002; Weerdesteyn et al. 2008). Most rehabilitation strategies intended to improve walking function report success in approximately 50% of cases (Nadeau et al. 2013; Awad et al. 2016). This limited success could be due in part to poor understanding of the specific neural circuitry involved, how impairment of these circuits affects walking function, and appropriate strategies to treat affected individuals. Here we investigated the use of a neurophysiologic marker, the long-latency reflex (LLR) response in the ankle musculature, to distinguish functional status among a heterogeneous sample of individuals with chronic stroke.

Muscle stretch produces both short- and long-latency reflex responses (Hammond 1955; Matthews 1970; Upton et al. 1971). Short-latency reflexes (SLRs) are mediated by the spinal cord (Pearson and Gordon 2013), while long-latency reflexes are mediated, at least in part, supraspinally (Cheney and Fetz 1984; Palmer and Ashby 1992). LLRs can be observed in both the upper (Deuschl et al. 1985; Day et al. 1991) and lower extremities (Brouwer and Ashby 1992; Nielsen and Petersen 1995; Petersen et al. 1998b). Evidence suggests that muscle stretch elicits a sequence of events: sensory pathways are activated, producing an ascending signal; sensory signals integrate information with cortical and subcortical structures; and ultimately an LLR response is produced in the homonymous muscle via descending motor pathways (Day et al. 1991; Petersen et al. 1998a). Because it involves communication between multiple cortical structures and can be modulated by changes in cortical excitability, this response is also known as the transcortical reflex (Phillips 1969; Scott et al. 2015).

Early studies focused on characterizing changes in upper extremity LLRs in neuropathologies (Lee and Tatton 1978; Matthews et al. 1990; Scott et al. 2015). A proportion of patients with a rare disorder known as Klippel-Feil syndrome have a bifurcation of corticospinal projections, causing LLRs to be present bilaterally following unilateral muscle stretch (Matthews et al. 1990). Parkinsonian rigidity is associated with an increase in LLR amplitude relative to healthy adults (Lee and Tatton 1975; Mortimer and Webster 1979). Some individuals with multiple sclerosis reveal diminished or even absent LLRs, which sometimes coincide with abnormal somatosensory or motor evoked responses (Deuschl et al. 1988; Michels et al. 1993). Diminished, or absent, LLRs have been described in some individuals following cortical stroke (Conrad and Aschoff 1977; Marsden et al. 1977b), while later onset and prolonged duration LLRs were revealed on the contralesional side in one individual with a right supplementary motor cortex lesion (Dick et al. 1987). In a sample of individuals with only thalamic stroke, LLRs were attenuated or absent, and these differences appeared to detect pathology better than somatosensory evoked potentials alone (Chen et al. 1998). While collectively these studies offered early insights concerning the presence of disordered physiology across multiple pathologies, they lacked information regarding how impaired LLRs relate to volitional motor function.

The LLR is known as the functional stretch reflex because it is present during volitional activity and is highly adaptable to varying biomechanical and task constraints (Jones and Watt 1971; Freedman et al. 1976; Scott et al. 2015). LLRs can be elicited and measured during either isolated dynamic ankle movements or coordinated movements like walking (Sinkjær et al. 1996; Petersen et al. 1998b). However, it is unclear whether the LLR plays an important role in steady-state walking. Observation that the LLR amplitude modulates in response to stretches imposed during the gait cycle led to the hypothesis that LLRs aid torque production at the ankle (Sinkjær et al. 1999). Because the LLR latency is shorter than volitional response times and the response can be modified by both central and peripheral conditions, the timing is ideal for monitoring and correction of modified task demands in response to environmental perturbations (Pruszynski and Scott 2012). Given the potential utility of LLRs to integrate information about both afferent and efferent systems, make early corrections, and aid in torque production, LLRs could be vital to successful walking. Furthermore, LLRs could serve as an important indicator of central nervous system functional status among individuals with chronic stroke. For example, the presence of intact LLRs may identify individuals better equipped to handle challenges encountered while walking, while diminished or absent LLRs may identify individuals who lack the same capacity for appropriate neuromotor adaptation.

With regard to walking mechanics following stroke, it is recognized that many individuals have weakness of the lower extremity musculature which is typically associated with propulsive power deficits (Lamontagne et al. 2002; Jonkers et al. 2009; Banks et al. 2017). The ankle joint power profile during walking typically reveals a large, distinct concentric peak of propulsive power during pre-swing, known as A2 (Winter 2009). A2 is mediated by the plantarflexors and generates both the potential and kinetic energy necessary to push off during late stance and transition the limb into swing (Winter 1983a). A2 is typically diminished following stroke; the extent to which A2 is decreased is strongly associated with walking speed, capacity for gait speed modulation, and severity of walking dysfunction (Winter 1983; Olney et al. 1991; Jonkers et al. 2009; Kitatani et al. 2016). Propulsive power generated by the plantarflexors also drives knee flexion during swing, one of the major contributors to limb clearance (Nadeau et al. 1999; Little et al. 2014). For all that is known about the biomechanical deficits following stroke, much less is known about the specific neural mechanisms underlying these impairments.

Here our goal was to assess the relationship between lower extremity LLRs, measures of supraspinal drive, and walking function following stroke. We characterized dorsiflexor LLRs during volitional ankle plantarflexion. We used transcranial magnetic stimulation (TMS) to assess how these LLRs relate to changes in supraspinal drive with varying task demands. Finally, we investigated the relationship between LLRs and clinical and biomechanical measures of lower extremity function in individuals with chronic stroke.

## 2. Methods

### 2.1. Subjects

Thirteen individuals with chronic stroke (age 63±8 yrs, 11 male) and 13 healthy age-matched (age 61±8 yrs, 6 male) controls participated. Demographic data for the stroke cohort, reported in Table 1, illustrate a diverse group of individuals with mild-to-moderate motor impairment. Study inclusion criteria for the stroke group included: a diagnosis of unilateral cortical or subcortical stroke at least six months prior to date of enrollment, ability to follow three-step commands, and ability to walk at least ten meters without assistance. Stroke diagnosis was confirmed by CT or MR imaging results in medical records. All participants were free of any contraindications for transcranial magnetic stimulation, including implanted metal above the chest, seizure disorders, or pregnancy (Rossini et al. 2015).

**Table 1.** Subject Demographics.

| Subject | Age (yrs) | Sex | Paretic Side | Chronicity (mos) | Stroke Type | Lesion Location | LE FMA | AFO/Device | LLR Status |
| --- | --- | --- | --- | --- | --- | --- | --- | --- | --- |
| Stroke 01 | 80 | M | R | 97 | Hemorrhagic | L temporal lobe | 32 | -- | LLR+ |
| Stroke 02 | 53 | M | R | 25 | Ischemic | L parietal lobe | 34 | -- | LLR+ |
| Stroke 03 | 61 | M | R | 35 | Ischemic | L ICA occlusion with striatocapsular involvement | 24 | Aircast® | LLR+ |
| Stroke 04 | 61 | M | R | 87 | Ischemic | L temporal, parietal, and frontal lobes | 12 | Custom AFO | LLR- |
| Stroke 05 | 73 | M | R | 62 | Ischemic | L parietal lobe | 34 | -- | LLR+ |
| Stroke 06 | 67 | M | R | 86 | Ischemic | L parietal with basal ganglia involvement | 34 | -- | LLR+ |
| Stroke 07 | 69 | M | L | 267 | Hemorrhagic | R internal capsule | 17 | Walk Aide® | LLR- |
| Stroke 08 | 55 | F | L | 25 | Ischemic | R temporal lobe | 34 | -- | LLR+ |
| Stroke 09 | 57 | M | R | 59 | Ischemic | L frontal and temporal lobes with basal ganglia involvement | 13 | Custom AFO | LLR- |
| Stroke 10 | 62 | M | L | 65 | Ischemic | R parietal and temporal lobes | 34 | -- | LLR+ |
| Stroke 11 | 56 | F | R | 131 | Ischemic | L occipital and parietal lobes | 19 | Custom AFO | LLR+ |
| Stroke 12 | 54 | M | L | 7 | Hemorrhagic | R putamen | 22 | Custom AFO | LLR- |
| Stroke 13 | 66 | M | R | 131 | Ischemic | L insula and basal ganglia | 34 | -- | LLR+ |

Abbreviations: yrs, years; M, male; F, female; L, left; R, right; mos, months; ICA, internal carotid artery; LE FMA, lower extremity Fugl-Meyer motor score; AFO, ankle foot orthosis; LLR, long-latency reflex. Additional subject characterization is included in Fig. 3.

Testing occurred at the Brain Rehabilitation Research Center in the Malcom Randall VA Medical Center in Gainesville, FL, and spanned 2-3 days for each participant. All procedures were approved by the University of Florida Health Science Center Institutional Review Board and all participants gave written informed consent prior to participation. Testing was conducted in accordance with the Declaration of Helsinki.

### 2.2. Instrumentation

Isolated plantarflexion movements were tested using a commercially available dynamometer (Biodex System 3.2, Shirley, NY) and controlled by a Power 1401 data acquisition system (Cambridge Electronic Design Limited, Cambridge, England). Surface electromyography (EMG) was collected using a commercially available system (MA300-28, Motion Lab Systems, Baton Rouge, LA) from the medial gastrocnemius (MG), soleus (SOL), and tibialis anterior (TA) muscles using gel surface electrodes (Cleartrace 2, ConMed, Utica, NY) and snap-on preamplifiers (MA-420, Motion Lab Systems, Baton Rouge, LA). Electrodes were placed according to SENIAM guidelines (Freriks et al. 1999). Single pulse transcranial magnetic stimulation was applied using a Magstim BiStim^2^ with a 110 mm double cone coil (Whitland, UK). A Brainsight TMS neuronagivation system (Rogue Resolutions Ltd, Cardiff, UK) was used to maintain coil placement.

Analog signals from the dynamometer (torque, position, and velocity) were lowpass filtered using an analog hardware filter (100 Hz cutoff). All data were recorded in Signal 6.0 (Cambridge Electronic Design Limited, Cambridge, England) at a sampling rate of 2000 Hz.

### 2.3. Protocol

Participants were tested on the dynamometer in a single session. Each participant was seated in the Biodex chair with the seatback fully upright and the paretic leg (or test leg in healthy controls) extended, with approximately 90 degrees of hip flexion, 20 degrees of knee flexion, and the ankle positioned against the foot plate at neutral plantar/dorsiflexion, as shown in Fig. 1a. All joints were positioned so movement would occur only through the sagittal plane at the ankle. This configuration minimizes contributions of the hip muscles to plantarflexion while simultaneously positioning the medial gastrocnemius and soleus muscles within the optimal operating ranges (Winter and Challis 2008; Rubenson et al. 2012). The test leg was randomized across all healthy control participants. Transcranial magnetic stimulation (TMS) was localized to generate maximal motor evoked responses (MEPs) in the MG and SOL. Resting motor threshold (rMT) was determined as the minimum stimulus level required to elicit a response ≥100 μV peak-to-peak amplitude in three out of five trials (Rothwell et al. 1999).

**Fig. 1.**
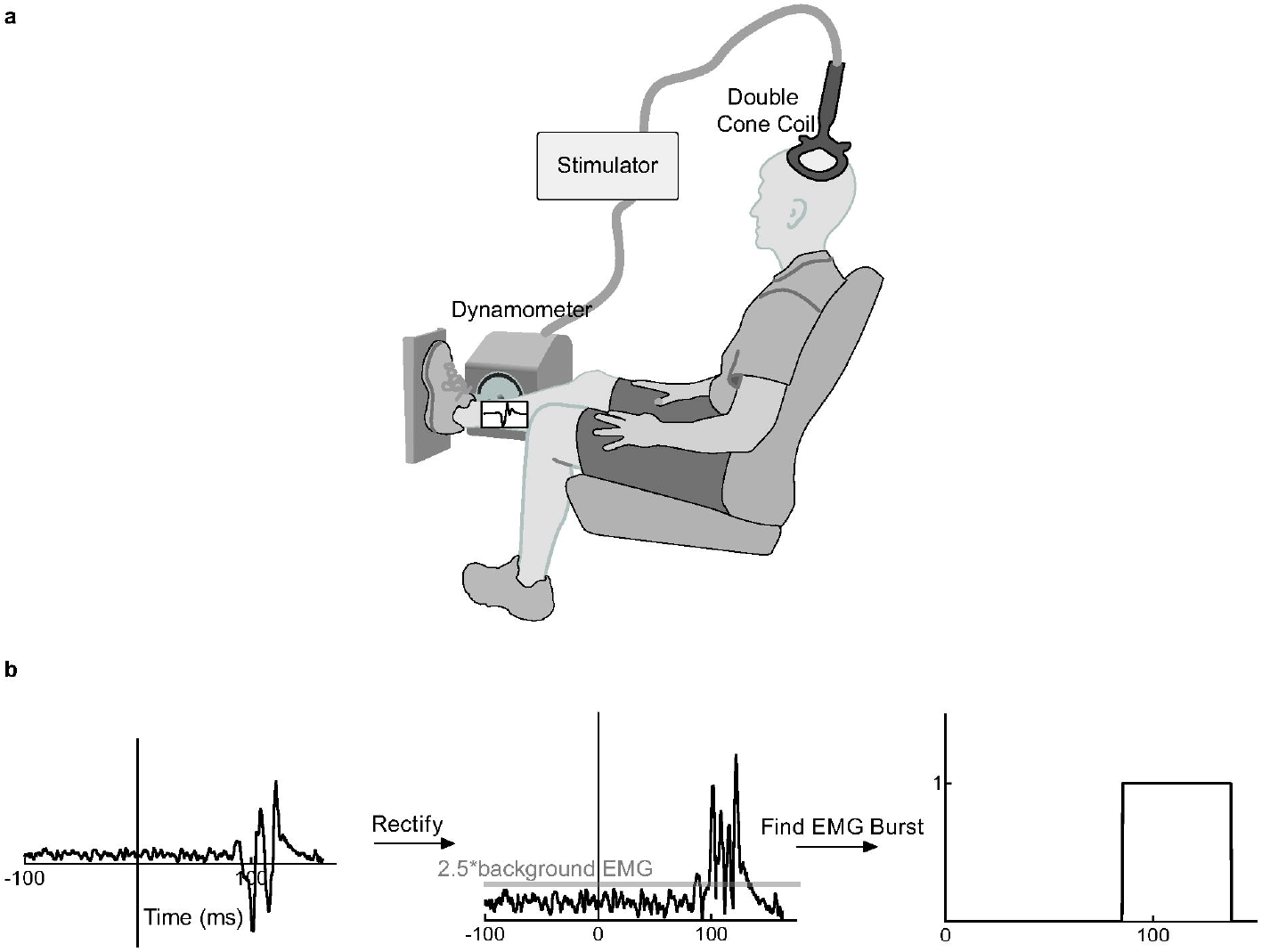
**a)** Illustration of experimental setup. Participant positioned with test leg extended and foot resting against dynamometer footplate, with ankle aligned to axis of rotation. Transcranial magnetic stimulation (TMS) was delivered through a double cone coil positioned above contralateral hemisphere motor cortex. **b)** LLR detection algorithm. Developed using healthy control data, LLRs were detected by rectifying the EMG and then searching for a burst that exceeds 2.5 times the background EMG prior to the onset of movement. Once this threshold is achieved for a minimum of 10 msec, the burst is transformed into a step function, allowing for consistent identification of LLRs.

During testing, participants were instructed to generate and hold 10-20% of their maximum voluntary contraction (MVC) torque against the dynamometer foot plate. Participants were provided real-time visual feedback of torque output to ensure consistent effort and task attention. Following a one second hold of 10-20% MVC torque, a magnetic stimulus was applied at 120% of SOL rMT. In the isometric condition, the foot plate remained stationary in the neutral position during stimulation. In the dynamic condition, the foot plate started in approximately five degrees of dorsiflexion. Following the one-second hold, the footplate released, allowing the participant to plantarflex, “as hard and as fast as possible,” up to a maximum velocity of 90 degrees per second through their available range of motion. This rate is comparable to, or slower than, angular velocities that occur at the ankle during walking. Magnetic stimulation was triggered when the ankle moved through the neutral position. After each trial, the participant was given a 2-3 second rest before the foot plate was returned to the starting position. A minimum of six trials were performed in each test condition. Some individuals were unable to achieve a sufficiently large range of motion prior to the arrival of the magnetic stimulus, causing a superposition of LLR and MEP responses that could not be separated, and were therefore excluded from this analysis.

### 2.4. Data analysis

Data were processed offline using Matlab R2015a (The MathWorks, Natick, MA). EMG was filtered using a 4^th^ order bandpass filter (10-450 Hz cutoff range). In the isometric condition, background EMG was measured 100 milliseconds prior to the magnetic stimulus to determine an activity threshold for each trial (mean ± 1 standard deviation). In the dynamic condition, the length of the activity threshold window was adjusted manually for each participant to include only the period prior to movement onset (range of 50-100 ms). This difference in establishing duration of the activity threshold window for the dynamic condition was to exclude long-latency reflex activity from the background EMG calculation.

### 2.5. Outcome measures

Primary outcome measures include LLR presence and TA MEP_area_ change. LLR presence was quantified as the percentage of trials in which the amplitude of an EMG burst within 100 - 170 milliseconds after movement initiation exceeded 2.5 times the average background EMG (Petersen et al. 1998b). Given that all controls showed LLRs, these criteria were determined using a detection algorithm developed on the healthy control data (Fig. 1b). LLR presence was measured only during dynamic plantarflexion. TA MEP_area_ is the area under the rectified and background-normalized motor evoked response elicited by TMS measured in the tibialis anterior muscle. We have expressed TA MEP_area_ change as the ratio between the isometric and dynamic conditions using the following equation:

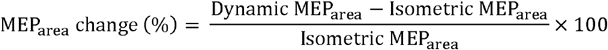

Secondary clinical outcomes for this analysis include: Lower Extremity Fugl-Meyer motor score, Short Physical Performance Battery (SPPB) score, Dynamic Gait Index (DGI), self-selected walking speed (SSWS), and fastest comfortable walking speed (FCWS). The Lower Extremity Fugl-Meyer Motor Function Score is a subscale of the Fugl-Meyer Assessment of Motor Recovery After Stroke, a widely used assessment of motor impairment with a maximum value of 34 points (Fugl-Meyer et al. 1975). The SPPB measures mobility and balance performance on a twelve-point scale and demonstrates robust predictive capacity for all-cause morbidity and mortality in older adults (Guralnik et al. 1994; Volpato et al. 2011). The DGI evaluates functional stability during walking with a maximum score of 24 points and is validated for use in chronic stroke (Shumway-Cook and Woollacott 1995; Jonsdottir and Cattaneo 2007). SSWS and FCWS were measured as the average speed from 3-5 passes over a 16-foot pressure-sensitive walkway (GaitRite Platinum Plus System, Version 3.9, Havertown, PA). For SSWS, participants walked at their casual, comfortable pace. FCWS was assessed as the fastest speed the participant could safely attain when walking, “as if you are crossing the street and the walk signal changes to a red hand.” Clinical measures were administered within a single session by a licensed physical therapist (VLL).

Our secondary biomechanical outcome for this analysis is peak ankle plantarflexor power (A2). A2, the second peak in the sagittal plane ankle power profile, corresponds to plantarflexor power generation (Winter 2009). A2 was derived from inverse dynamics using motion analysis performed while participants walked at their self-selected speed on an instrumented split-belt treadmill (Bertec, Columbus, OH). Marker data were obtained with a 12-camera Vicon motion capture system (Vicon MX, Vicon Motion Systems Ltd., Oxford, UK) using a modified Helen Hayes marker set sampled at 200 Hz. One healthy control did not complete the instrumented gait assessment. Ankle power data during gait are not available for one stroke participant due to dependence on a rigid ankle foot orthosis during gait assessments. All other participants walked on the treadmill with either an Aircast^®^ or without a brace and produced valid kinetics.

### 2.6. Statistical analysis

Data were assessed for normality using the Shapiro-Wilk W test and were not normally distributed (p’s < 0.05). Therefore, TA MEP_area_ change and walking speeds were assessed for group differences using Kruskal-Wallis ANOVA and a significance level of α = 0.05. Post-hoc analyses were carried out using the Steel-Dwass method to correct for multiple comparisons. Clinical assessments were compared between subgroups of individuals stratified by LLR presence using Mann-Whitney U tests. A Bonferroni correction was applied to these comparisons, and significance assessed using α = 0.017. One-tailed Spearman correlations assessed the relationships between MEP_area_ change and A2, using α = 0.05. All tests were carried out in JMP Pro 11 (SAS Institute, Cary, NC).

## 3. Results

All thirteen healthy controls revealed long-latency reflexes in response to rapid stretch of the TA during voluntary plantarflexion. Nine individuals post-stroke, herein referred to as LLR present (LLR+), also showed this stretch-mediated EMG. Four individuals post-stroke, referred to herein as (LLR-), lacked long-latency EMG activity in response to TA stretch. These three patterns are exemplified in Fig. 2a. Twenty of the twenty-six individuals tested showed facilitation of TA MEP_area_ in the dynamic, relative to the isometric, condition (Fig. 2b). There was a significant effect of group (p = 0.047) on MEP_area_ change. The post-hoc tests revealed significant facilitation of TA MEP_area_ (p = 0.029) in LLR- individuals, with a median (IQR) of 251% (130-404) relative to controls, with a median of 49% (−10-108).

**Fig. 2.**
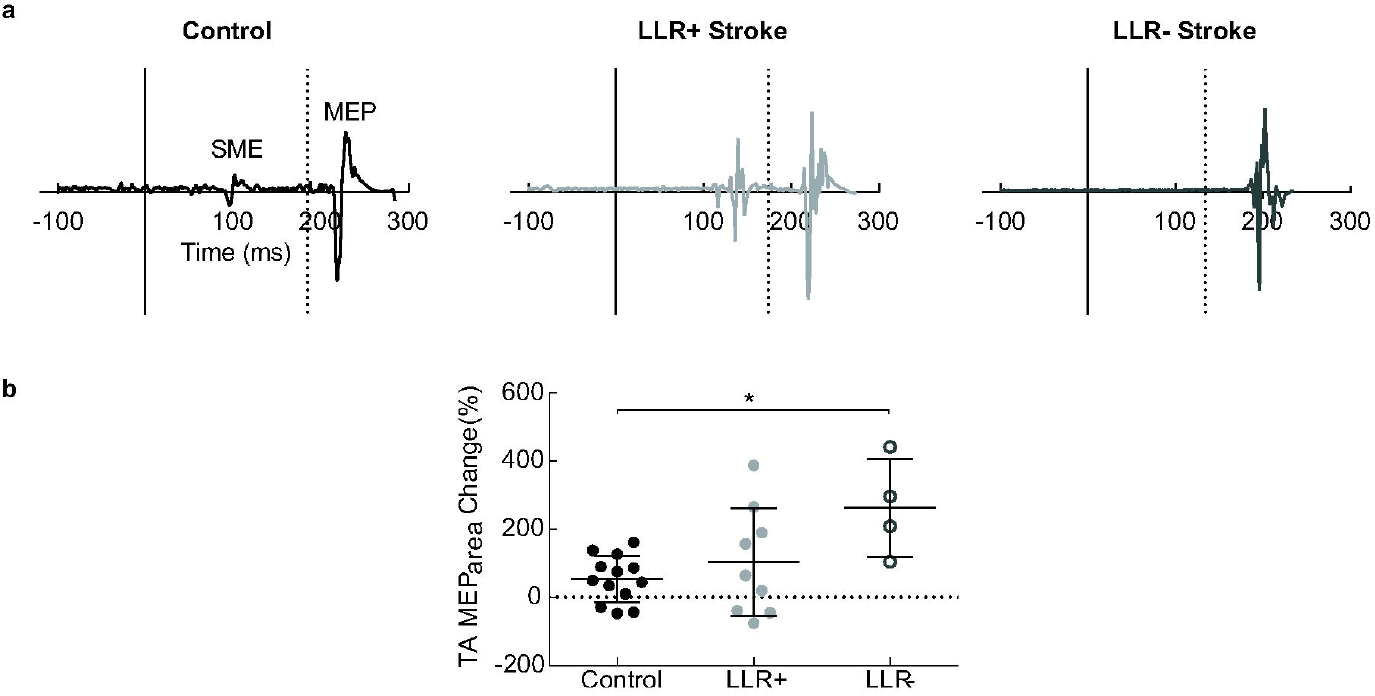
Long-latency reflex responses are absent in some individuals post-stroke. **a)** Left, example healthy control EMG shows long-latency reflex (LLR) activity with a latency of approximately 100 milliseconds (ms) from movement onset (time=0), followed by transcranial magnetic stimulation pulse (TMS, dotted line) and motor evoked potential (MEP) beginning at approximately 220 ms. Center, example EMG for one LLR+ individual post-stroke, showing LLR and the later MEP. Right, example EMG for an individual post-stroke who does not exhibit LLR (LLR-). **b)** LLR- individuals show exaggerated MEP_area_ change relative to healthy controls. * indicates significance at p<0.05.

LLR- individuals revealed lower clinical scores and walking speeds than LLR+ and healthy controls (Fig. 3a). Fugl-Meyer motor score was lower in LLR- individuals, with a median (IQR) score of 15 (12.25— 20.75), than LLR+ individuals with a median score of 34 (28—34; p = 0.007). SPPB score was also lower in LLR- individuals, with a median score of 5.5 (5—6.75), than LLR+ individuals with a median score of 11 (9—12; p = 0.008). DGI score did not show a significant difference between LLR- individuals when considering the multiple comparisons correction, with a median score of 10 (8.5—13), and LLR+ individuals with a median score of 22 (18.5—23.5; p = 0.02). Kruskal-Wallis ANOVA detected differences across groups for SSWS (p = 0.0004) and FCWS (p = 0.001, Fig. 3b). Post-hoc analyses revealed that SSWS was higher in healthy controls, with a median speed of 1.3 m/s (1.2—1.5), than both LLR+, with a median speed of 1.1 m/s (0.96—1.2; p = 0.01), and LLR-, with a median speed of 0.32 m/s (0.22—0.66; p = 0.01), and higher in LLR+ than LLR- (p = 0.03). FCWS was higher in healthy controls, with a median speed of 2.0 m/s (1.7—2.3) than LLR+, with a median speed of 1.6m/s (1.2—1.9, p = 0.03), and LLR-, with a median speed of 0.48m/s (0.34—0.95; p = 0.01), but was not significantly different between LLR+ and LLR-(p > 0.05).

**Fig. 3.**
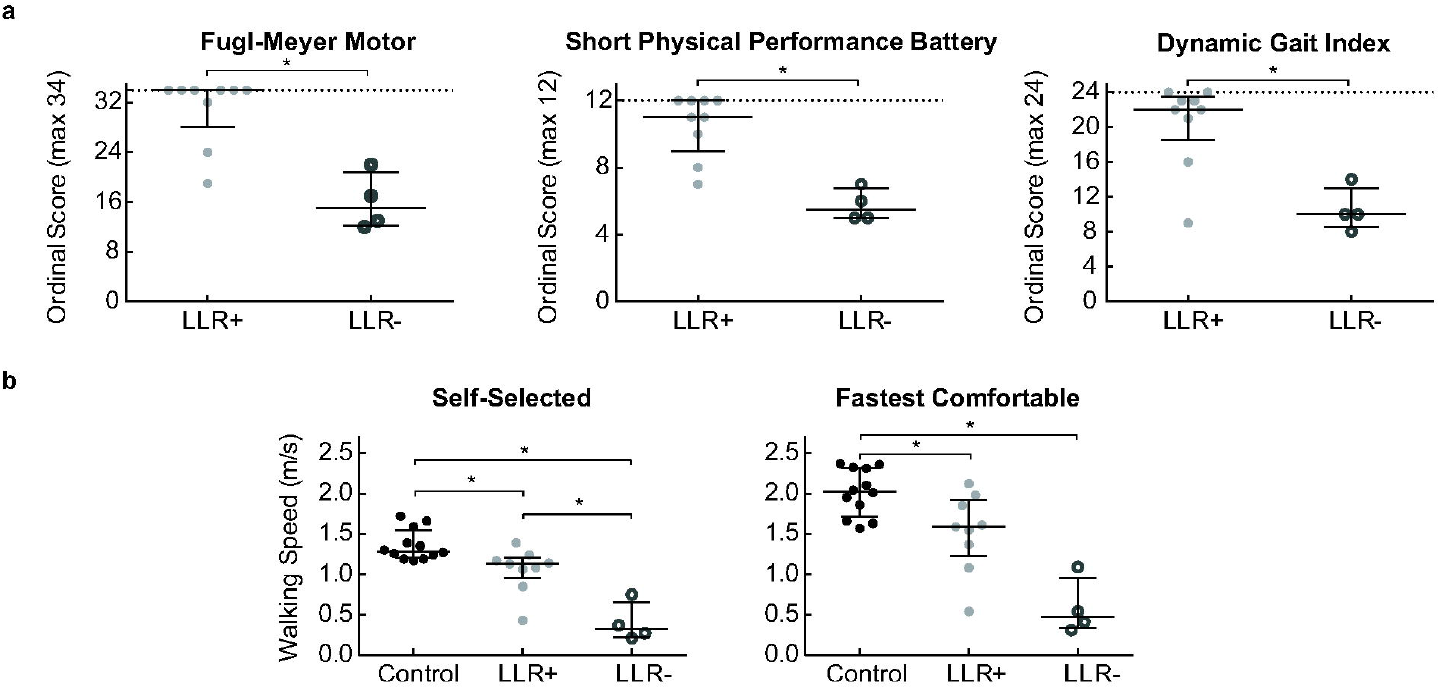
**a)** Clinical scores are markedly lower for LLR- individuals than LLR+ individuals post-stroke. From left to right, scores include: Fugl-Meyer lower extremity motor assessment, Short Physical Performance Battery, and Dynamic Gait Index. **b)** Walking speed was different between all groups for self-selected walking speed, left, and between controls and each LLR group for fastest comfortable walking speed, right. Bars represent median ± interquartile range. * indicates significance at p<0.017 for clinical scores or p<0.05 for walking speeds.

Tibialis anterior MEP_area_ change was not associated with A2 magnitude in healthy controls (p = 0.14, Fig. 4), however, the stroke group revealed a significant correlation (p = 0.01). The scatterplot in Figure 4 illustrates an unambiguous gap between low and high ankle power; of note, all LLR- individuals produce low ankle power.

**Fig. 4.**
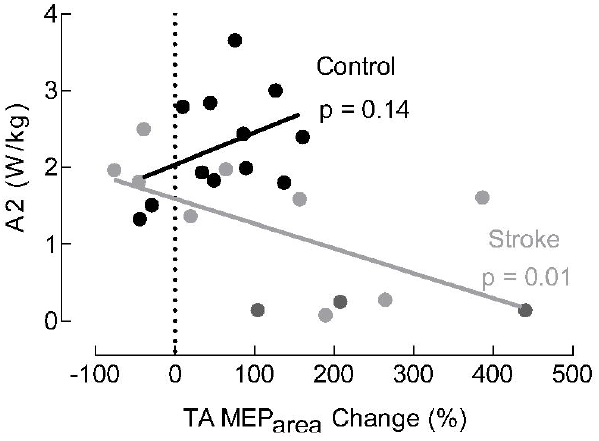
A MEP_area_ change predicts ankle plantarflexor power (A2) post-stroke, but not in healthy individuals. In healthy controls, the two variables are not correlated (Spearman ρ=0.34, p>0.05 black), however in individuals post-stroke there is a significant negative correlation between MEP_area_ change and A2 (ρ=−0.64, p=0.03). LLR+ individuals are indicated in light gray circles, while LLR- individuals are indicated with gray open circles, for illustrative purposes.

## 4. Discussion

Our primary findings are dysregulation of TA MEPs during voluntary, dynamic plantarflexion in a subset of individuals post-stroke, a portion of whom also lack long-latency stretch responses in the TA during the same task. These two observations indicate physiological dysfunction that appears to be related to functional motor deficits in the lower extremity post-stroke. The lowest functioning individuals in our sample lack LLRs and exhibit profound dysregulation of antagonist MEPs during dynamic plantarflexion. Absence of LLRs likely signals dysfunction of the transcortical reflex pathway, while facilitated antagonist MEPs may indicate dysregulation of the reciprocal inhibitory circuit.

In this study, nearly one-third of individuals post-stroke lacked long-latency activity in the tibialis anterior during voluntary plantarflexion. While the resultant number of LLR- individuals is small and creates unbalanced groups for statistical comparisons, we posit that this proportion may be maintained within a larger sample. This stratification according to physiological criteria revealed unambiguous differentiation of functional status among these individuals. In contrast, characterization solely on the basis of clinical scores revealed heterogeneous groupings. Some of the individuals assessed did not have a sufficiently large range of motion available at the ankle to elicit stretch responses prior to the arrival of the MEP and were therefore excluded from this sample. One participant had limited dorsiflexion range and was instead positioned at neutral prior to movement in the dynamic condition, and is included in this analysis. Such methodological challenges limit our ability to draw definitive conclusions about the prevalence of LLR- individuals among the population living with chronic stroke, but we are able to discuss the characteristics of those individuals that lacked LLRs within the context of this experiment.

This observation of LLR- individuals confirms early research conducted in the upper extremity in stroke and other pathologies involving supraspinal lesions (Conrad and Aschoff 1977; Marsden et al. 1977b; Lee and Tatton 1978; Deuschl et al. 1988). The long-latency reflex has a strong cortical component that integrates sensory inputs from multiple pathways and creates functionally relevant output (Jones and Watt 1971; Freedman et al. 1976; Palmer and Ashby 1992). It is not surprising, then, that individuals with cortical and subcortical lesions could have disrupted transcortical reflex pathways. Stretch reflexes of all latencies are important for maintenance of steady-state walking, obstacle negotiation, and responses to perturbations (Dietz et al. 1984; Sinkjær et al. 1996). As a functional motor correlate, absence of the LLR could contribute to lower clinical scores, slower walking speeds, and more impaired gait mechanics. A significant limitation of current clinical assessments is that they are mere descriptors, providing only an approximation of behavior at the individual level. For example, two individuals with similar lower extremity Fugl-Meyer scores can have quite different clinical and biomechanical presentations. The underlying pathophysiology in these same individuals likely differs as well. Elucidating the neural mechanisms contributing to behavioral manifestations of motor impairment requires a deeper understanding of the neurophysiologic differences between high and low functioning individuals.

Early evidence suggested potential diagnostic significance to long-latency reflexes, but this line of research was largely abandoned prior to its completion (Lee and Tatton 1978). In a sample of individuals with multiple sclerosis, LLRs were able to detect pathology in a manner complementary to, yet sometimes distinct from, other non-invasive assessments such as motor- and somatosensory evoked potentials (Michels et al. 1993). Due to the nature of the transcortical reflex, the somatosensory tracts must be functioning appropriately in order to elicit an LLR (Marsden et al. 1977a); the same would be expected from the motor system. The relationship between LLR dysfunction and clinical and functional measures could relate to the proposed roles of the transcortical reflex in postural stability and corrective responses (Evarts and Fromm 1981). Combined with the work of others, our findings suggest that the LLR may be a sensitive indicator of pathology within this sensorimotor pathway following stroke (Lee and Tatton 1978). Importantly, LLRs are more straightforward to assess than other evoked potentials that require specialized and expensive laboratory equipment. Although they may only contribute a portion of the information necessary to build a complete understanding of an individual’s condition, LLRs could represent an essential tool for identifying individuals with pathology in motor control mechanisms of the lower extremity (Deuschl et al. 1988).

The underlying mechanism responsible for the exaggerated facilitation of TA MEPs remains unclear. Given the dynamic condition instructions, it is not unreasonable that the controls exhibited a small facilitation in TA MEP size due to a generalized increase in motor excitability during volitional plantarflexion. But the excessive facilitation present in some individuals post-stroke warrants further consideration. Diminished reciprocal inhibition, and even a reversal pattern termed reciprocal facilitation, have been observed in some neuropathologies including: cerebral palsy, spinal cord injury, and stroke (Gottlieb et al. 1982; Myklebust et al. 1982; Crone et al. 2003). The appearance of reflex reversals is inconsistent and poorly understood, but may be attributable to the disynaptic reciprocal inhibitory circuit (Okuma and Lee 1996; Crone et al. 2003; Bhagchandani and Schindler-Ivens 2012). In addition to spinal circuitry, supraspinal inputs to inhibitory interneurons contribute to the reciprocal inhibitory pathway (Lundberg 1970). Lesions in the motor cortex may, therefore, interrupt normal patterns of inhibitory control, allowing for pathologic disinhibition with dynamic movement. Our observation that the magnitude of TA MEP facilitation during plantarflexion is negatively correlated with the magnitude of ankle plantarflexor power may be indicative of over-excitable dorsiflexor activity, inhibiting plantarflexion vital to gait. The mechanism of this dysregulation warrants further investigation. The finding that all LLR- individuals and some LLR+ individuals within this sample exhibited excessive facilitation indicates that this facilitation is likely driven by a different mechanism than the LLR; the interaction between these two responses would require further investigation within a larger sample.

In combination, long-latency reflexes, supraspinal drive as measured by TMS, and clinical and functional measures offer a comprehensive assessment of neurophysiologic and behavioral function after stroke. To our knowledge, this is the first time that LLRs have been investigated in parallel with walking function in chronic stroke. We chose to assess LLRs during isolated plantarflexion because many individuals with chronic stroke lack the capacity to activate the plantarflexors during walking, limiting their ability to generate enough propulsive force to walk efficiently (Marks and Hirschberg 1958; Jonkers et al. 2009; Banks et al. 2017). It is also challenging to accurately assess walking after stroke because many individuals cannot walk completely unassisted (i.e., without a brace, cane, or weight support) for long durations, all of which is required for thorough biomechanical analysis (Jorgensen et al. 1995). Because neurophysiology is a contributor to observable behavior, there is a need to systematically investigate both neurophysiology and behavior in tandem in order to understand disordered motor control. Insights gained from assessing these pathophysiologic patterns can contribute to understanding the heterogeneity among individuals with chronic stroke. Additionally, this approach may enable identification of individuals who possess enough neural substrate for rehabilitation interventions to be effective versus those who may not. If this is the case, LLRs could serve as a biomarker for recovery of sensorimotor function following stroke. Biomarker development requires determination of analytical validity, qualification, and utilization (Institute of Medicine Committee on Qualification of Biomarkers and Surrogate Endpoints in Chronic Disease 2010). Future research is required to determine the appropriateness of LLRs as a biomarker within this population, as well as a more acute population of individuals post-stroke. Lower extremity LLR dysfunction could either be related to the stroke itself or a consequence of maladaptive plasticity that develops over time, but this cannot be discerned from assessing only individuals in the chronic phase.

The goal of the present study was to assess the relationship between the presence of lower extremity LLRs, modulation of supraspinal drive, and walking function in chronic stroke. A notable subset of individuals with stroke lacked a normal physiologic response to a physiologically relevant muscle stretch. These same individuals exhibited dysregulated motor evoked responses in the tibialis anterior during active plantarflexion and were the lowest-functioning participants within our study. Such individuals are likely to have different rehabilitation needs than their higher functioning counterparts. Future work is necessary to identify the major contributors to LLR dysfunction within these individuals, assess the mutability of these responses and the transcortical reflex pathway, and determine the functional consequences.

## Conflict of Interest Statement

The authors declare that they have no conflict of interest.

## Acknowledgements

This work was supported by the Department of Veterans Affairs, Rehabilitation RR&D Service [Grant #01435-P and Research Career Scientist Award #N9274S, Patten, PI], Portions of this project were presented in abstract form at the 2016 American Physical Therapy Association Combined Sections Meeting in Anaheim, CA and the 2016 Biomechanics and Neural Control of Movement Conference in Mt. Sterling, OH. The authors thank Spencer Gilleon for assistance with data collection and Theresa McGuirk, M.S., Jennifer Bromwell, and Janke Mains-Mason for assistance with data processing.

